# Neurofeedback training for improving motor performance in healthy adults: A systematic review and meta-analysis

**DOI:** 10.1101/2022.04.26.487963

**Authors:** Ryoji Onagawa, Yoshihito Muraoka, Nobuhiro Hagura, Mitsuaki Takemi

## Abstract

Neurofeedback training (NFT) refers to a training where the participants voluntarily aim to manipulate their own brain activity using the sensory feedback abstracted from their brain activity. NFT has attracted attention in the field of motor learning for its potential to become an alternative or additional training method for general physical training. In this study, a systematic review of NFT studies for motor performance improvements in healthy adults and a meta-analysis on the effectiveness of NFT were conducted. To identify relevant studies published between January 1st, 1990 to August 3rd, 2021, a computerized search was performed using the databases, Web of Science, Scopus, PubMed, JDreamIII, and Ichushi-Web. Thirty-two studies were identified for the qualitative synthesis and 13 randomized controlled trials (286 subjects) for the meta-analysis. The meta-analysis revealed significant effects of NFT for motor performance improvement examined at the timing after the last NFT session (standardized mean difference = 0.96, 95% CI = 0.40–1.53), but with the existence of publication biases and substantial heterogeneity among the trials. Subsequent subgroup meta-analysis demonstrated reliable benefits when the NFT is performed longer than 1 week. The effectiveness of NFT for each motor performance measurement (e.g., speed, accuracy, and hand dexterity) remains unclear because of high heterogeneity or due to small sample size. Further accumulation of empirical NFT studies for motor performance improvement will be necessary to provide reliable evidence about the NFT effects on specific motor skills and to safely incorporate NFT into real-world scenarios.

## Introduction

A skill to fluently perform actions is not only important in particular competitive settings (e.g., sports), but it is also a building block for our daily behavior. There is an inherent link between the state of brain activity and the performance of the associated motor tasks. For instance, the cortical oscillatory activity of the beta frequency (13–30 Hz) is implicated to be involved in movement preparation, generation, and termination (Kilavik et al., 2013), such that the modulation of beta-power correlates with reaction times (He et al., 2020; Senkowski et al., 2006). Furthermore, the higher power of sensorimotor rhythm (SMR) preceding the task is associated with better performance in motor precision tasks (e.g., shooting skills) (Cheng et al., 2017). If such evidence indicates an existence of an optimal brain state for higher motor task performance, training to make the brain activity influenced toward that brain state may be beneficial to improve motor performance.

Neurofeedback training (NFT) is a training method to regulate one’s own brain activity toward the targeted brain state by using feedback based on the features of the ongoing brain activity. A previous study indicated that reinforcing participants’ performance by modulating the brain activity towards the target state can indeed lead to an improvement in task performance (Shibata et al., 2011). The effectiveness of NFT for treating diseases and disorders has been reported, such as in the domain of motor rehabilitation following a stroke (Wang et al., 2018), posttraumatic stress disorder (Chiba et al., 2019; Koizumi et al., 2016; van der Kolk et al., 2016), major depression (Fernández-Alvarez et al., 2022; Trambaiolli et al., 2021), and attention deficit hyperactivity disorder (Arns et al., 2009; Cortese et al., 2016; Enriquez-Geppert et al., 2019; Lofthouse et al., 2012; Micoulaud-Franchi et al., 2014). Some studies also target healthy individuals to improve motor performance (Gong et al., 2020; Mirifar et al., 2017; Xiang et al., 2018), executive function (Viviani and Vallesi, 2021), and memory accuracy (Yeh et al., 2021). However, to accurately estimate the effect of NFT, collective evaluation of the reports in the field as a whole is necessary, rather than individual reports. In this paper, we perform a systematic review and a meta-analysis, focusing on the effectiveness of NFT on improving motor performance.

Meta-analyses of NFT on motor function have been conducted. Several studies have examined whether NFT can improve motor performance in sports athletes (Mirifar et al., 2017; Xiang et al., 2018). Moreover, the effect of NFT on motor rehabilitation following a stroke has been examined (Bai et al., 2020). In both analyses, some benefit of NFT has been shown, but the generalizability of evidence in a general healthy population remains unclear. Recent technological advances (i.e., easily accessible brain activity measurement devices) have accelerated the public use of NFT. Therefore, it could be beneficial to evaluate the effectiveness and reliability of NFT in various motor tasks in a nonspecialized group of participants.

In this study, a systematic review of NFT studies for motor performance improvements in healthy adults and a meta-analysis on the effectiveness of NFT were conducted. In the meta-analysis, detailed effects of NFT relative to the following categorical items were evaluated in subgroup analyses. First, the effects of NFT with three types of control conditions were compared: sham neurofeedback, general training without neurofeedback, and conditions without any intervention. Comparison with the sham condition, where incorrect or mock feedback is used, helps eliminate the placebo effect and determine the actual effect of the NFT. Comparisons with general training protocols such as physical training, mental imagery, and meditation will provide practical insights. NFTs are useful only when the effect level is comparable with those of the non-NFT. Second, a subgroup meta-analysis was conducted to examine the NFT effect on different motor tasks. The extent to which the effects were moderated by the participants’ demographics, brain imaging modalities, and intervention periods were also examined.

## Methods

The systematic review and meta-analysis were conducted following the Minds Handbook for Clinical Practice Guideline Development 2020 ver 3.0 chapter 4 (Minds Manual Developing Committee, 2021).

### Database search

Searches of PubMed, Scopus, Web of Science, JDreamIII (Japanese scientific manuscript database), and Ichu-shi (Japanese medical manuscript database) were conducted to identify articles published in English and Japanese from January 1, 1990 to August 9, 2021. Two authors (RO and YM) independently conducted the database search. The following terms were used for the search. The terms of brain imaging modalities included “electroencephalogram” (EEG), “magnetoencephalogram” (MEG), “functional magnetic resonance imaging” (fMRI), “functional near-infrared spectroscopy” (fNIRS), and “positron emission tomography” with both abbreviated and nonabbreviated forms. For neurofeedback related terms, “neurofeedback,” “neural feedback,” “biofeedback,” “brain-computer interface” (also with abbreviated form), “brain-machine interface” (also with abbreviated form), “real-time,” and “feedback” were used. Motor performance related terms included “motor,” “sensorimotor,” “movement,” “sport,” “athlete,” “muscle,” “skill,” “performance,” “ability,” “strength,” “accuracy,” “reaction time,” “response time,” ““speed,” “learning,” “training,” and “control”. For the search, the asterisk function in the search engines was used (e.g., “muscl*”) to ensure that all the related articles were captured. The above terms in Japanese were used to search the Japanese databases. Within each set of terms (i.e., brain imaging, neurofeedback, and motor performance-related terms), the terms were combined with the “or” function on search engines. The three “or”-combined sets of terms were then combined with the “and” function. Furthermore, a manual search was conducted through the reference sections of the identified articles to detect studies that were dismissed from the database search.

### Study selection

Articles were screened in two phases. The first screening was performed to remove obviously irrelevant articles on the basis of the relevance of titles and abstracts. The second screening was performed by checking the full article with respect to the eligibility criteria (see below). The first screening was independently performed by RO and YM. Studies with disagreements between the two individuals were passed to the second screening. RO and YM again independently performed the second screening, and the other two authors (NH and MT) conducted a full-text assessment of the studies that were disagreed upon. The four reviewers discussed the articles until a consensus was reached.

Studies included for the qualitative synthesis satisfied all of the following eligibility criteria: (i) The participants were healthy adults with the age range of 18-65; (ii) noninvasive brain imaging methods were used; (iii) NFT was performed; (iv) motor performance was measured as an outcome. Here, any observable muscular outputs as motor performance was considered, which includes the speed of the reaction times, precision and dexterity of movements, and the balance of the whole body; (v) a control group was set for the NFT intervention; (vi) it was an original article published in a scholarly peer-reviewed journals and written either in English or in Japanese; (vii) the study design was either a randomized controlled trial (RCT) or a non-randomized controlled study (NRS). In addition to these criteria, studies included for the meta-analysis satisfied (viii) the outcome information is enough to calculate the effect size of the intervention; (ix) the study design was RCT; and (x) the study was assessed as a low or moderate risk of bias (see below in the section of quality assessment).

### Data extraction

RO and YM individually extracted the following data during the second screening: study design, participant’s demographics, types of intervention and control conditions, and outcome measures regarding behavioral motor performance and the type of brain activity. For studies included in the qualitative synthesis, RO and YM further extracted details about interventions and experimental protocols. Moreover, descriptions of serious adverse events and side effects were recorded.

For the meta-analysis, motor performance measures (statistical values) after the last NFT session from RCTs were extracted. Here, each experiment in each paper was regarded as a single trial. If the study was a crossover design, only the results of the first intervention were extracted, and those following the second intervention were not considered for the analysis. This is to avoid possible carryover effects from the first intervention. For the studies using factorial design, the results of the single condition intervention groups were used. For each trial (i.e., data from each experiment), motor performance data were extracted in the form of means, standard deviations, and the number of participants in each group. The extracted data were filled out on a predesigned spreadsheet that automatically calculates the standardized mean difference (SMD) as the effect size. We did not contact the authors to request the missing outcome data or to ask for more detailed information about the study.

### Data analyses

A systematic review of studies identified after the screening process was conducted. First, the study population (i.e., sample size, age, sex, and history of sports experience), NFT protocols (i.e., brain imaging modalities, types of cortical activity used for the NFT, the amount of intervention, type of feedback, and type of control), and other variables (i.e., year of publication, study design, and outcome measure) were analyzed. For the meta-analysis, a standard pairwise meta-analysis was conducted using a random-effects model by combining all data of the same outcome into a single dataset to avoid selective analysis of outcomes. Publication bias was visualized as a funnel plot and was estimated via Begg’s test and Egger’s test. The publication bias was evaluated only if there were more than 10 studies for the particular target outcome. If significant publication bias was detected (*p* < 0.1 was considered statistically significant for both tests), the trim-and-fill analysis was conducted to correct for the publication bias.

The heterogeneity was evaluated by estimating the variance between studies (χ^2^ test and I2 statistic). Subsequently, a subgroup meta-analysis was conducted using Cochrane’s Q test with a mixed-effects model to account for the between-subgroup heterogeneity. The individual alpha level was adjusted using the Bonferroni–Holm method to control the overall experiment-wise error rate. Analyses were conducted for the following categorical items: (i) types of control conditions, categorized as the general training, sham feedback, or no intervention; (ii) participant’s demographics, categorized as athletes or nonathletes; (iii) brain imaging modalities used in NFT, categorized as electric (e.g., EEG and MEG) or metabolic (e.g., fNIRS and fMRI); (iv) targeted motor performance, such as movement accuracy (e.g., golf putting and archery shooting), reaction times, and manual hand dexterity; (v) intervention period, categorized as longer or shorter than 1 week. Furthermore, a meta-regression analysis was conducted to test the relationship between the effect size and the total number of NFT sessions and the sum of NFT time.

The meta-analysis, subgroup analysis, and related visualizations were conducted using Review Manager version 5.4. Begg’s test, Egger’s test, trim-and-fill analysis, and funnel plot visualizations were conducted by using the metafor package in R version 4.1.1. The meta-regression analysis was conducted using the meta package in R version 4.1.1.

### Quality assessment

RO and YM independently conducted the risk of bias assessment for the studies in which the statistics of motor performance measures were extracted. The assessment was conducted using the Cochrane Collaboration’s tool (Higgins et al., 2011). Low, high, or unclear risk of bias was determined for each of the following six domains: (i) selection bias, determined by random allocation sequence, baseline imbalance, and allocation concealment; (ii) performance bias, determined by blinding of participants and personnel; (iii) detection bias, determined by blinding of outcome assessments; (iv) attrition bias, determined by the incompleteness of outcome data and intention-to-treat analysis; (v) reporting bias, determined by selective reporting; (vi) other possible sources of bias, such as conflict of interest (COI), a sample size determination method, and incorrect statistical methods. Disagreements were resolved through discussions by the four authors. Individual studies judged at a high risk of bias in more than two domains were excluded from the meta-analysis.

## Results

### Search results

Figure 1 shows the literature review process. We obtained 2307 articles through databases and 18 articles from the reference sections of the retrieved studies. Of these, 2214 articles were excluded due to the title and the abstract irrelevance. One hundred and eleven articles passed to the full-text screening for the full-text eligibility assessment. Here, 79 articles were excluded on the basis of the following reasons: research using patients (n = 3), research using adolescents (n = 2), review articles (n = 2), no NFT (n = 16), study without a control group (n = 16), and an outcome measurement that did not include motor performance (n = 40). Full texts from the 32 remaining studies were retained for the qualitative synthesis. Finally, 11/32 RCT studies were eligible for the meta-analysis. The reasons for the exclusion were as follows: the reported results were insufficient to calculate effect sizes (n = 12), the studies were non-RCTs (n = 7), and the studies had a high risk of bias (n = 2).

**Figure 1.**
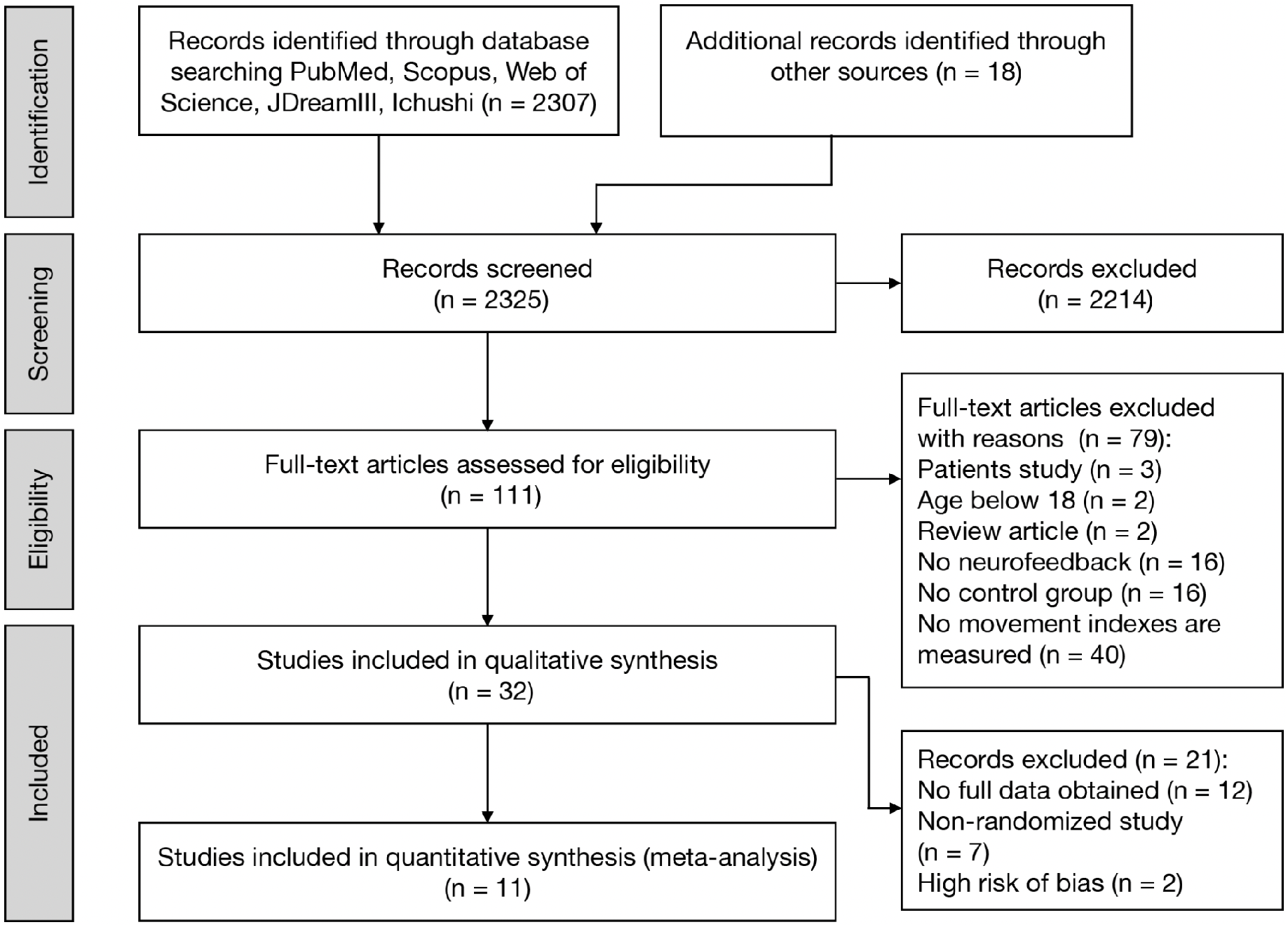
PRISMA flowchart of literature search and inclusion.

### Study and sample characteristics

Table 1 summarizes the main characteristics of 32 studies. The publication years of these studies were between 1991 and 2021 (Fig. 2A). More than half of them were published in the years after 2017. Fourteen studies focused on athletes, and one study focused on music students.

**Figure 2.**
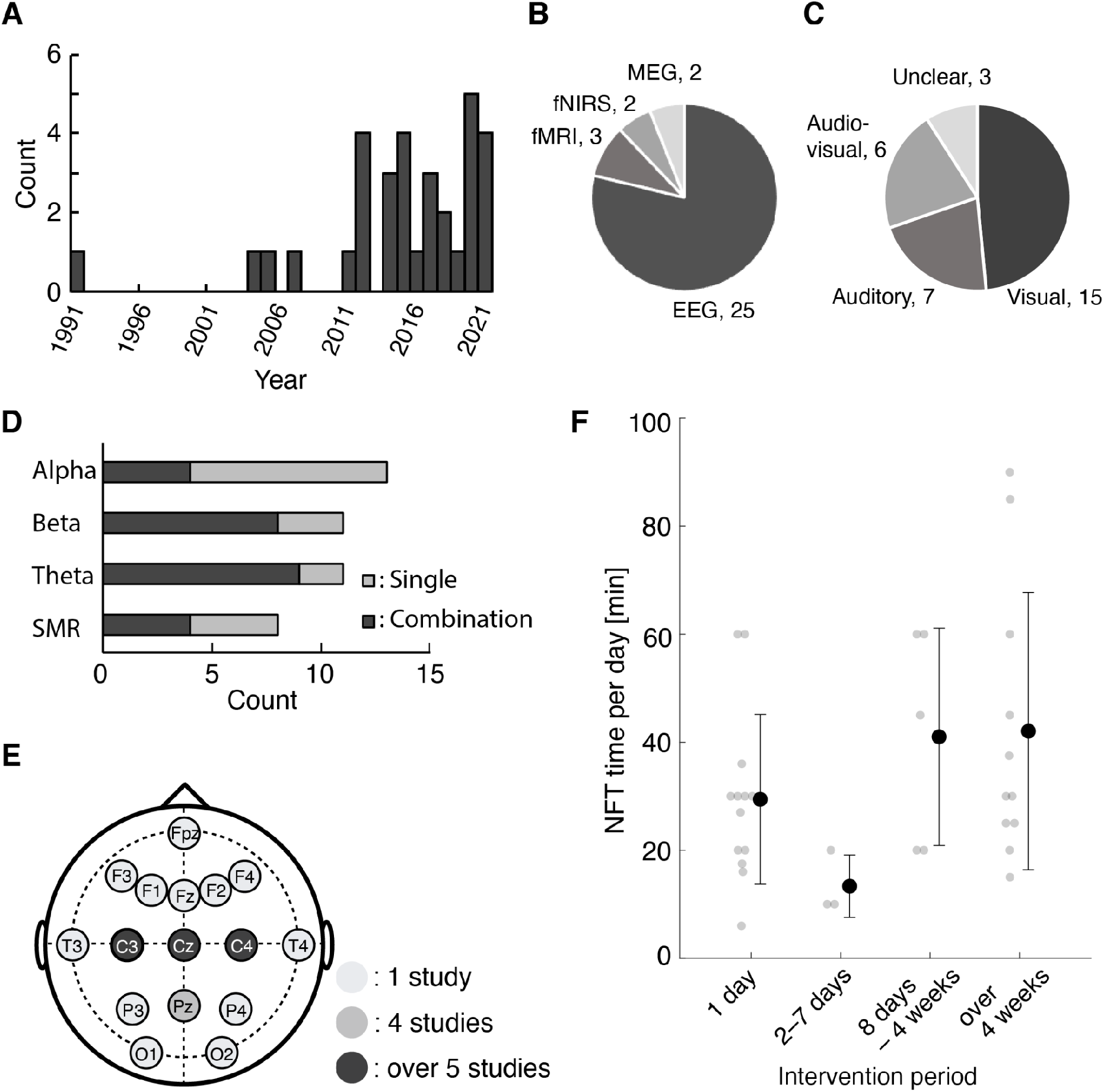
Results of the qualitative assessment. (A) Yearly publications. (B) Brain imaging techniques and (C) feedback modalities of brain activity used for NFT. (D) The proportion of EEG frequency bands and (E) EEG recording channels utilized across 25 EEG-based NFT studies. (F) Daily NFT duration grouped by the length of the intervention period. Gray dots represent the results of each trial. Black dots and error bars represent the mean and the standard deviation of each group.

**Table 1.**
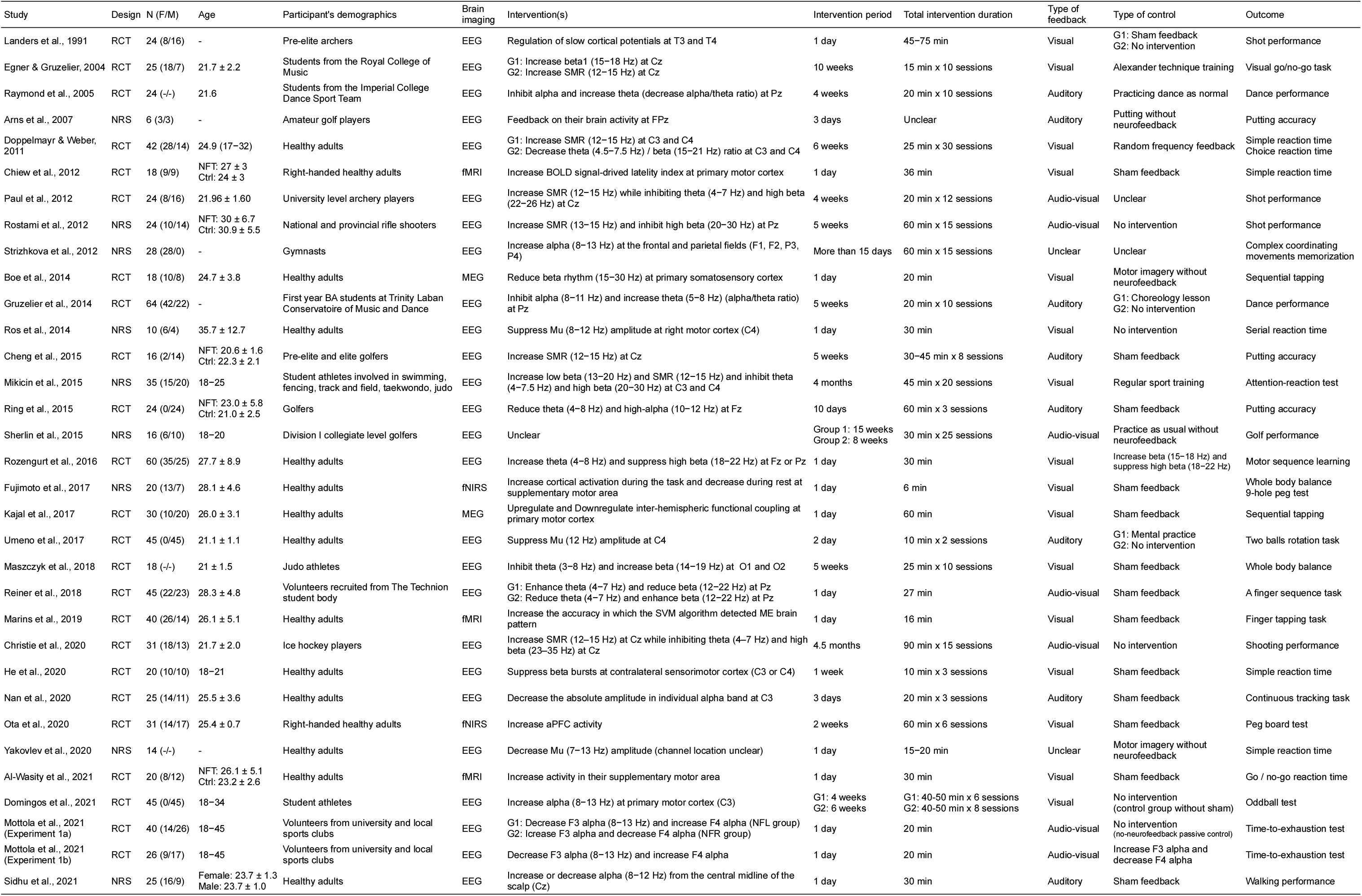
Experimental characteristics of the studies included in the qualitative synthesis.

The brain imaging modalities used for neurofeedback were as follows: 25 studies were EEG, two studies were MEG, three studies were fMRI, and two studies were fNIRS (Fig. 2B). The sensory modality of the feedback, which was transformed from the targeted brain activity, was either visual, auditory, or audio-visual (Fig. 2C). For EEG-based NFT, the frequency bands used in the neurofeedback were theta (4–7.5 Hz), alpha (8–13 Hz), sensorimotor rhythm (SMR) (12–15 Hz), and beta (15–30 Hz) (Fig. 2D), and electrodes around the sensorimotor cortex (C3, Cz, C4) were most frequently used (12 studies; Fig. 2E). Alpha rhythm and SMR were often used alone, and theta and beta rhythm tended to be used in combination (e.g., theta/beta ratio). Intervention periods ranged from 1 day to 4.5 months, with daily NFT duration ranging from 6 to 90 min (Fig. 2F). For the control conditions, 14 studies used the sham feedback, three studies used other neurofeedback protocols (i.e., different EEG frequency bands), nine studies conducted general training without neurofeedback (e.g., mental practice), and eight studies set the control condition as a condition without intervention (i.e., participants maintained at rest without doing anything).

For motor performance, eight studies focused on hand dexterity measured with a finger tapping task, nine-hole peg test, or pegboard test. Eight studies focused on movement accuracies such as shot performance or golf putting skills. Seven studies focused on speed factors measured with simple reaction time, go/no-go task, or choice reaction time.

### Risk of bias assessment

None of the studies (13 RCTs) satisfied the criterion to be judged as low risk of bias for all assessed domains (Fig. 3). For the domain of selection bias, most studies did not report how randomization was performed for assigning participants to each group. Moreover, 4 studies did not match the baseline level of motor performance between the groups prior to the NFT. For the allocation concealment, only two studies that reported the details of an allocation method were assessed as low risk, and the other studies had an unclear risk. For the performance bias, 10 studies were assessed as high risk because of insufficient blinding of experimental information to participants or personnel. For the detection bias, all studies were assessed as having unclear risks because of the lack of blinding information to those assessing the outcomes. For the attrition bias, one study that reduced the number of participants without any reasons was judged as a high risk. For the reporting bias, none of the studies have pre-registered the protocol. Furthermore, three studies that did not describe how the brain activity changed after the NFT were assessed as high risks. Since the assumption of NFT is that the training can induce the brain activity change, lack of such description makes it difficult for us to evaluate the underlying cause of the motor performance changes by the NFT, if any. Finally, for other possible sources of bias, two studies that had the potential of using inappropriate statistical tests were classified as high risks, and seven studies that did not report either COI or the method for determining sample size were assessed as having unclear risks. Overall, two studies that were judged as having high risks for more than two domains were excluded from the meta-analysis.

**Figure 3.**
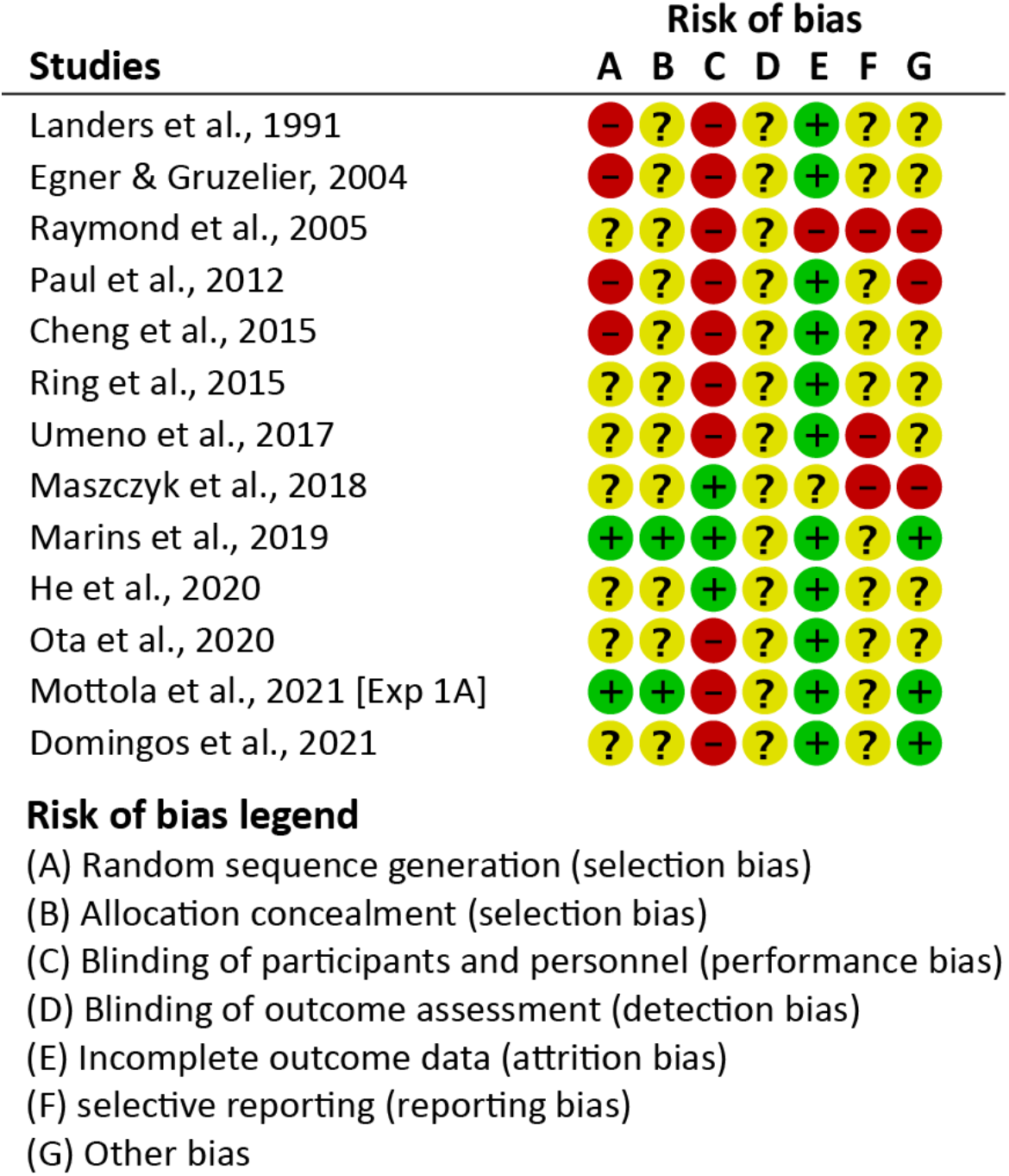
Risk of bias assessment. Green, low risk of bias; red, high risk of bias; yellow, unclear risk of bias.

### Overall effects of NFT for improving motor performance

Thirteen trials (total sample size: 286 participants) from 11 studies were used for the meta-analysis. Overall, NFT showed moderate to large effects on motor performance (SMD = 0.96, 95% confidence interval (CI) [0.40–1.53], Z = 3.32, *p* < 0.001; Fig. 4). However, a significant asymmetry was found in the funnel plot (Begg’s test: tau = 0.64, *p* = 0.002; Egger’s test: Z = 2.99, *p* = 0.003), indicating the existence of publication biases. For the trim-and-fill analysis, two artificial samples were imputed to adjust for potential biases (Fig. 5). The adjusted effect size was 0.62 (95% CI [-0.11-1.34], Z = 1.67, *p* = 0.09).

**Figure 4.**
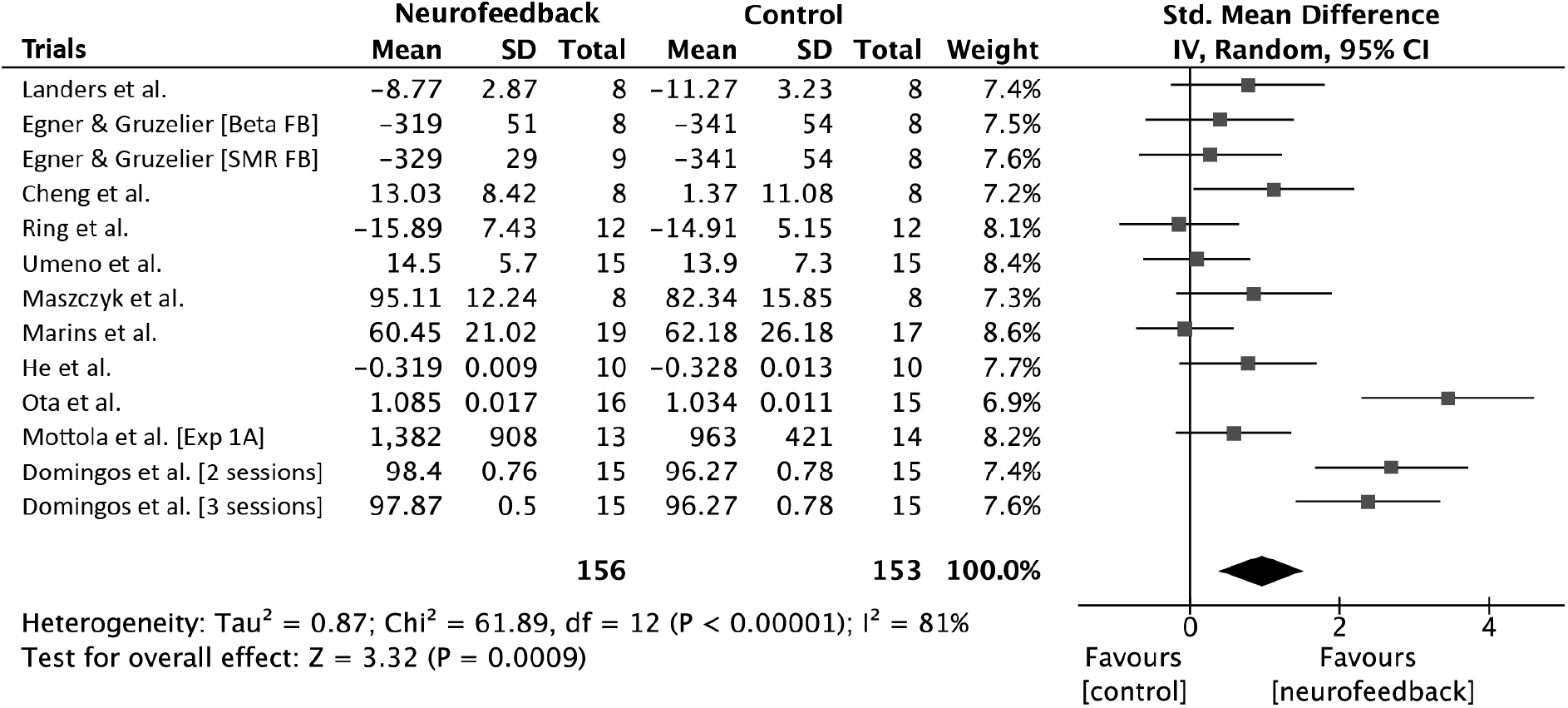
Forest plot for the efficacy of NFT in motor performances. The plot depicts all effect size estimates (SMD) and a 95% CI.

**Figure 5.**
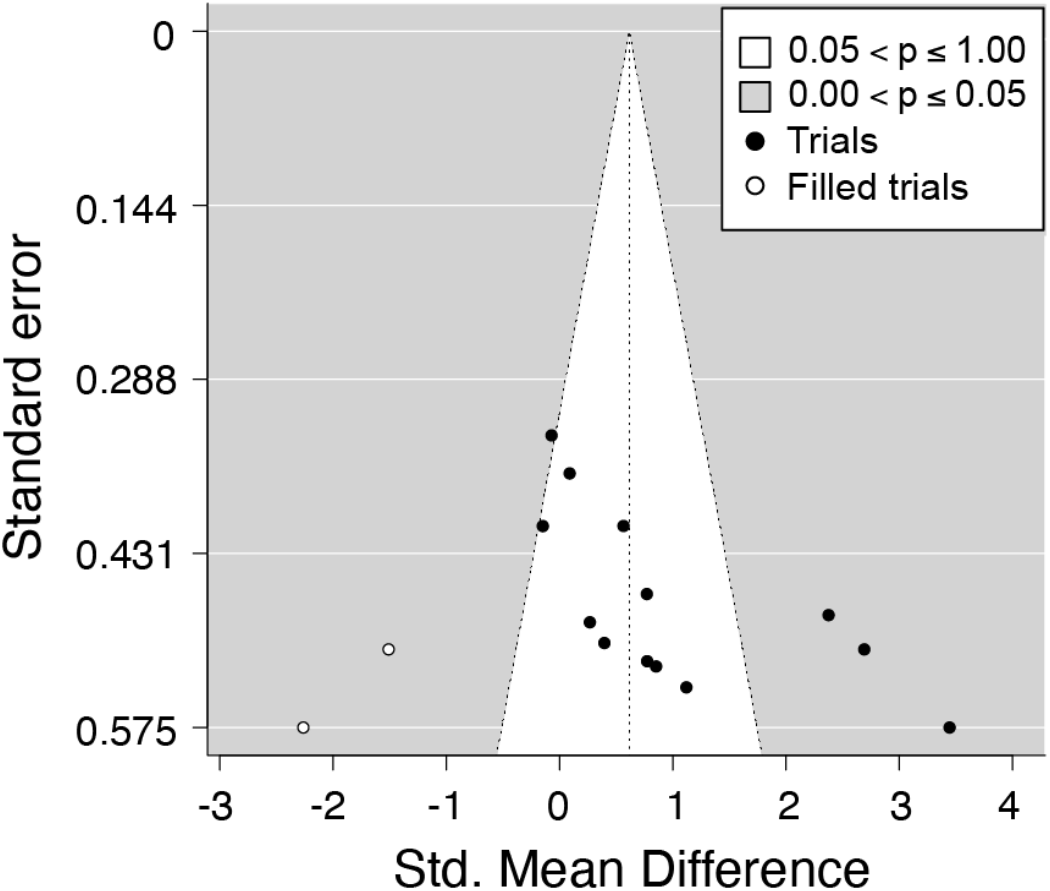
Funnel plot displaying effect sizes from the experimental results (solid circles) and imputed effect sizes calculated from the trim-and-fill analysis (open circles).

### Subgroup meta-analyses

There was a high level of heterogeneity between the studies (I^2^ = 81%). Hence, subgroup meta-analyses were conducted for the five categorical items: effects relative to control condition type, difference in the brain imaging modality, participant’s demographics, type of motor performance measure, and the duration of the intervention (Fig. 6).

**Figure 6.**
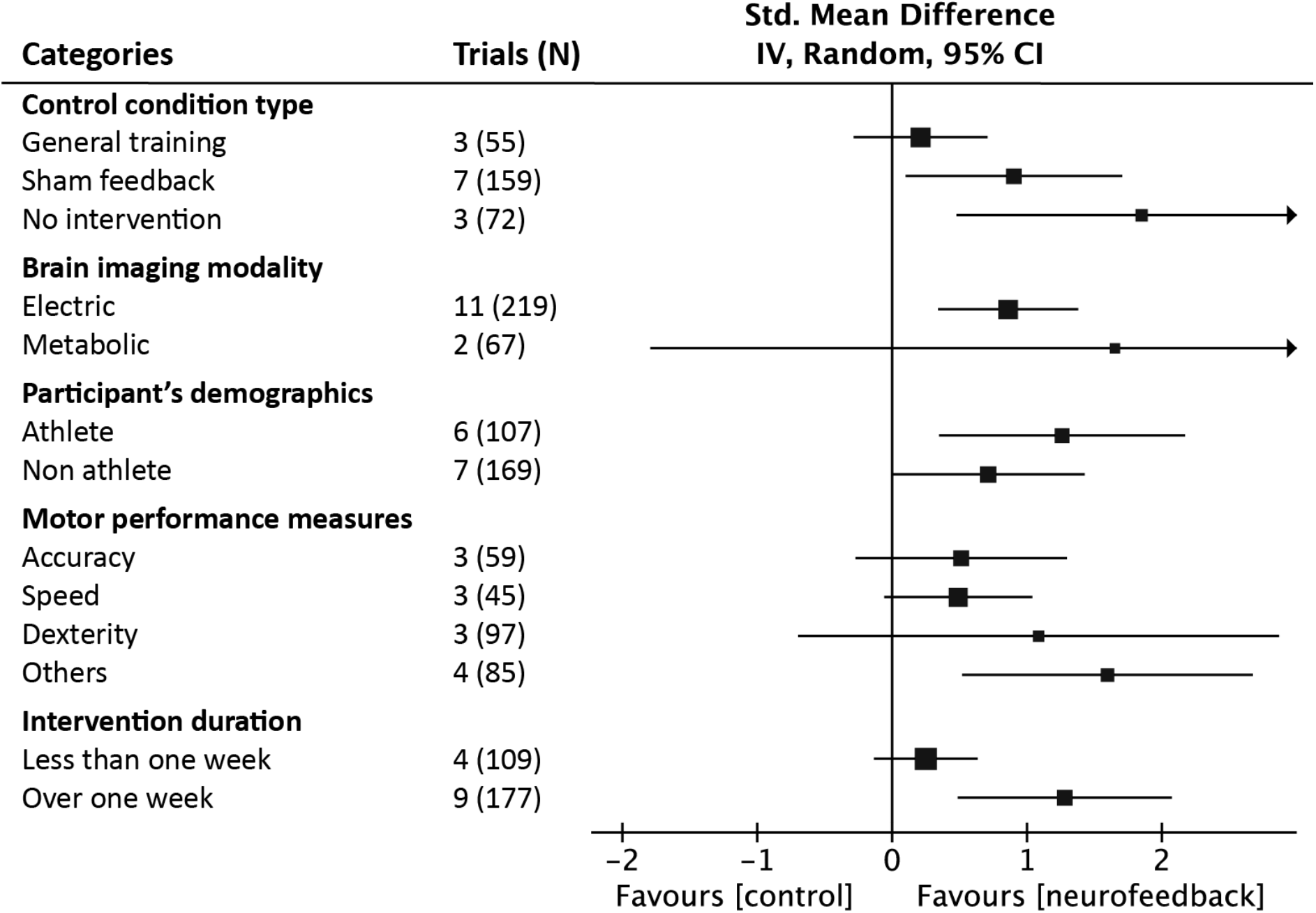
Results of subgroup analyses. The mean and 95% CI effect sizes were separately shown for control condition type, brain imaging modality, participants’ demographics, motor performance measures, and intervention period.

#### Effects relative to control condition type

We compared the effects of NFT with three types of control conditions: sham neurofeedback, general training without neurofeedback, and conditions without any intervention. The results showed a trend of subgroup differences for NFT (χ^2^ = 5.94, *p* = 0.05). Specifically, significant NFT effects were detected when comparing the conditions without any intervention (SMD = 1.85, 95% CI [0.48-3.22], Z = 2.64, *p* = 0.008; heterogeneity I^2^ = 85%) and sham feedback (SMD = 0.90, 95% CI [0.10–1.71], Z = 2.20, *p* = 0.03; heterogeneity I^2^ = 81%). However, NFT did not improve motor performance significantly better than general training (SMD = 0.21, 95% CI [-0.28-0.71], Z = 0.84, *p* = 0.40; heterogeneity I^2^ = 0%).

#### Effects of difference in brain imaging modality

The NFT trials were categorized into the ones using electric (EEG and MEG) or metabolic (fMRI and fNIRS) brain activity signals. The difference of NFT effects between the subgroups was not significant (χ^2^ = 0.20, *p* = 0.65). The effect of NFT in the electric group was medium to large (SMD = 0.86, 95% CI [0.33-1.38], Z = 3.24, *p* = 0.001), and the heterogeneity level was moderate (I^2^ = 67%). The effect of NFT in the metabolic group was not significant (SMD = 1.65, 95% CI [-1.79-5.10], Z = 0.94, *p* = 0.35; heterogeneity I^2^ = 96%).

#### Effects of participant’s demographics

In the participant’s demographics analysis, the trials were categorized into athlete studies or nonathlete studies. The subgroup difference was not significant (χ^2^ = 0.86, *p* = 0.35). The effect of NFT was medium to large for both the athletes (SMD = 1.26, 95% CI [0.35-2.17], Z = 2.72, *p* = 0.007) and nonathletes (SMD = 0.71, 95% CI [0.00-1.43], Z = 1.95, *p* = 0.05). Strong heterogeneity was detected in both groups (Athletes: I^2^ = 81%; Nonathletes: I^2^ = 80%).

#### Effects of difference in the motor performance measures

For the outcome measures of motor performance, the trials were divided into 4 categories: accuracy, speed, dexterity, and others. The subgroup differences were not significant (χ^2^ = 3.73, *p* = 0.29). The effects of NFT on movement accuracy, speed, and hand dexterity were not significant (Accuracy: SMD = 0.51, 95% CI [-0.27-1.30], Z = 1.28, *p* = 0.20; Speed: SMD = 0.49, 95% CI [-0.06-1.04], Z = 1.74, *p* = 0.08; Dexterity: SMD = 1.09, 95% CI [-0.70-2.87], Z = 1.19, *p* = 0.23), and the heterogeneity was medium in accuracy and strong in dexterity (Accuracy: I^2^ = 50%; Speed: I^2^ = 0%; Dexterity: I^2^ = 93%). There was a small to large effect on other motor skills (SMD = 1.60, 95% CI [0.54-2.67], Z = 2.96, *p* = 0.003) along with high heterogeneity (I^2^ = 80%).

#### Effects of intervention duration

In terms of the intervention duration, the trials were divided into two categories: NFT intervention shorter than 1 week (four trials) or longer than 1 week (nine trials). This 1 week threshold was determined on the basis of the intervention duration, which appeared to have two distributions (Fig. 2F). There was a significant difference between the two intervention duration categories (χ^2^ = 5.33, *p* = 0.02). For interventions of less than 1 week, the effect of NFT was not significant (SMD = 0.24, 95% CI [-0.14-0.63], Z = 1.25, *p* = 0.21; heterogeneity I^2^ = 0%). By contrast, for interventions of more than 1 week, the effect of NFT was moderate to large, though the significant heterogeneity was observed (SMD = 1.28, 95% CI [0.49-2.07], Z = 3.18, *p* = 0.001; heterogeneity I^2^ = 83%).

### Meta-regression analysis

The result of the subgroup meta-analysis examining the effect of intervention duration implies the relationship between the dose of NFT and the degree of motor performance improvements. Therefore, a univariate meta-regression analysis was conducted, with the NFT effect as the dependent variable and the total number of training sessions or the cumulative training time (minutes) as predictor variables. The results showed that the number of sessions was not associated with the NFT effect (F_1,11_ = 0.71, *p* = 0.42; Fig. 7A). However, cumulative training time was significantly associated with the NFT effect (F_1,11_ = 11.35, *p* = 0.006; Fig. 7B), such that the training time (dose) explained 59.8% of the motor performance improvement.

**Figure 7.**
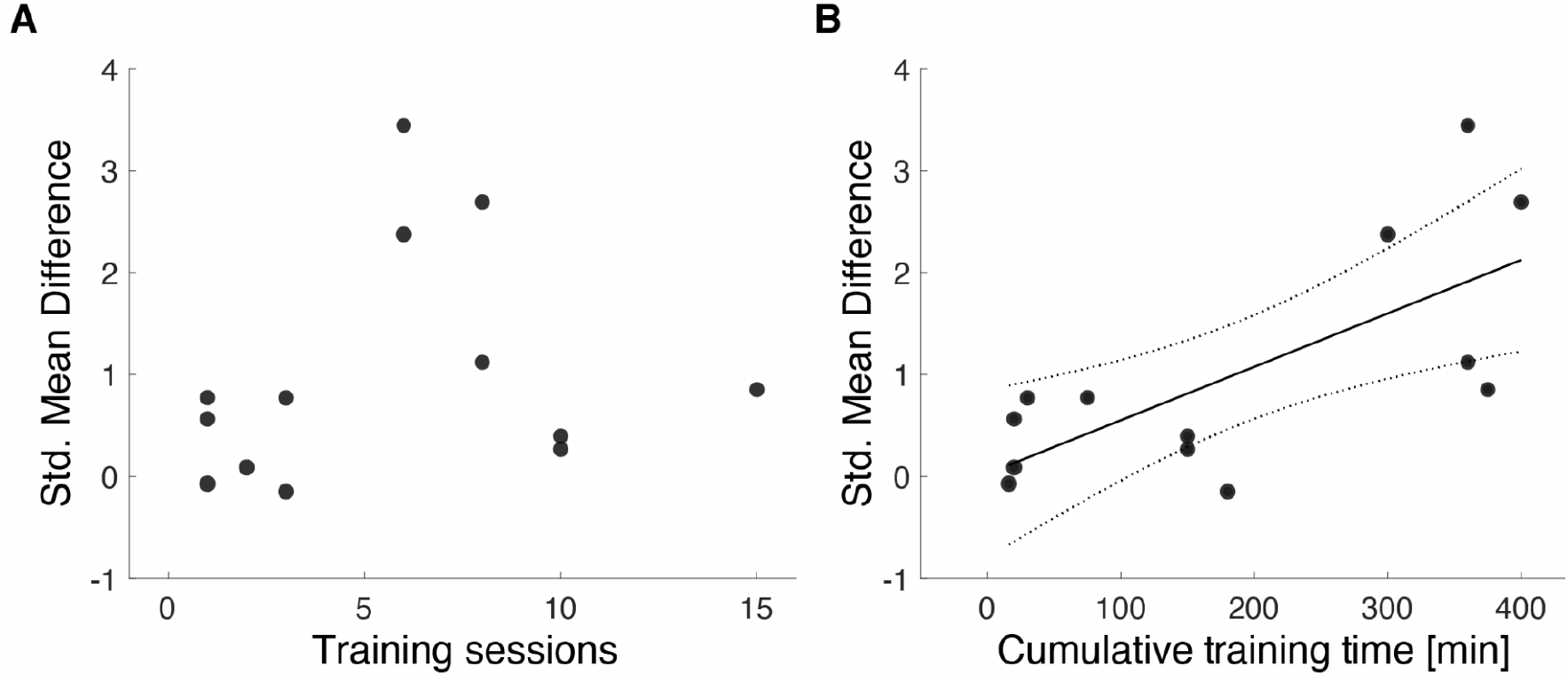
Relationship between the effects of NFT and (A) the total number of NFT sessions and (B) the sum of NFT time. Each dot represents the results from each trial. The regression line and its 95% CIs were presented only if the regression was statistically significant.

### Adverse effects

Side effects of NFT were evaluated. Only 5/32 studies mentioned the side effects. These five studies did not report any adverse effects during NFTs. Three studies assessed sleepiness, fatigue, or frustration using questionnaires. There was no difference in these side effects between the NFT and control groups.

## Discussion

This study examined the effect of NFT on motor performances by conducting a systematic review and meta-analysis. The effect on healthy adults was assessed, which differs from the previously conducted meta-analyses targeting athletes and patient populations. After the screening process, 32 studies were used for qualitative synthesis and 13 RCTs for the meta-analysis. Out of 32 studies, more than half were published later than 2017. This publication trend indicates that the field is still at its growing stage, and the data for the evaluation is still limited. Reflecting such a tendency, most of the studies were exploratory scientific research rather than a rigorously designed study to examine the effects of NFT. More evidence will likely be accumulated in the near future, which will increase the quality of the meta-analyses.

Although there was a significant effect of NFT on motor performance, correcting for the publication bias attenuated the effect. Therefore, overall, there was no conclusive evidence to support the benefit of NFT on motor performance. The publication bias may be diminished by an increase in sample size, which will lead to a more conclusive answer to the question in the future.

The results of the subgroup analysis suggested that the amount of intervention was related to the effects of NFT on motor performance. The NFT effect was greater in trials with an intervention duration of more than 1 week than with less than 1 week (Fig. 2F). Furthermore, a significant positive correlation was detected between the cumulative training time and the effect of NFT. These results suggest that a dose-response gradient exists in the NFT of motor performance. Since these findings are from exploratory analyses, future studies must clarify the function of this relationship, linear or sigmoidal, by using samples continuously covering the intervention durations. How long the effect will last after the final intervention would also be a target of interest for the future.

Similar to any other treatment, the effect of NFT could not be free from the placebo effect (Hammond, 2011). Therefore, comparing the NFT effect with a sham control group, not the group without any interventions, is necessary to evaluate the true effect of NFT. Indeed in the current meta-analysis, the effect size was smaller when the NFT was compared with its corresponding sham feedback group. This is likely because of the placebo effect (Cortese et al., 2016; Xiang et al., 2018), suggesting that a comparison with the sham feedback group is essential to assess the pure effects of NFT.

The effect of NFT was comparable with the effect obtained by general task training, indicating the possibility of NFT being a substitute for general motor training. However, the sample comparing the effects of NFT with the general training was small (three studies, 55 participants), preventing us from deriving any concrete conclusions. To assess the practicality of the NFT, it will be essential to examine the effectiveness of NFT with other training methods.

Although significant effects were identified in trials with intervention durations greater than 1 week, no significant improvements in outcome measurements for each subgroup (speed, accuracy, and dexterity) were observed. More data are certainly needed on each of the outcome measures to clarify the types of motor performances that can be expected to benefit from NFT. It is also worthwhile to examine the effects of NFT on outcomes that have not been examined currently (e.g., muscle power).

Brain activity that serves as the source signal for sensory feedback differs depending on the targeted motor performance. For instance, the neurofeedback system for the reaction time tasks has used EEG beta waves from the fronto-parietal region at rest or just before the onset of movements (Egner and Gruzelier, 2004; He et al., 2020), which have been shown to correlate with reaction times (Little et al., 2019; Tzagarakis et al., 2010). An NFT study aimed at improving putting accuracy has required participants to increase the SMR recorded at Cz (Cheng et al., 2015). Therefore, the neuronal mechanism of performance improvement by NFT is assumed to be different for each outcome. Future meta-analyses should increase the sample variability of both motor functions (e.g., speed, accuracy, power, and dexterity) and brain activity features (e.g., beta power and theta/beta ratio). It will be crucial to investigate which components of the motor task can be efficiently improved by the NFT using a particular brain activity pattern.

To use NFT for motor performance improvement, an assessment of NFT safety is necessary. In the current systematic review, only 5/32 included studies mentioned any description of the side effects. These five studies did not report any serious adverse effects during NFTs. This pattern is somewhat similar to the pattern observed in a systematic review examining the effect of NFT with cancer survivors (Luctkar-Flude and Groll, 2015). In this review, they found that only 4/17 studies mentioned the side effects, which were fatigue, sleep problems, headache, and a temporary intensification of previously problematic symptoms. NFT may cause transient side effects, although these side effects may not be due to the NFT protocol per se. They are rather general issues regarding any brain measurement procedures. Indeed, an RCT studying side effects associated with NFT has suggested that the side effects might be limited to tiredness and mild headaches (Rogel et al., 2015). We hope that future studies will at least describe the existence of side effects when performing the NFT so that a proper evaluation of NFT can be performed.

The assessment of bias risk highlighted numerous issues, resulting in a reduction in the certainty of the evidence. Common to most studies was the lack of implementation and reporting in that “specific methods of randomization and allocation of groups,” “how conditions were blinded,” “the number of subjects recruited, participated, data obtained, and analyzed,” and “the extent to which the analyzed measures and analytic methods were planned in advance.” In medicine, the effect of intervention is established by the collective contribution of individual studies, appropriately preregistering the study design and reporting the results on the basis of the reporting guideline such as CONSORT 2010 (Schulz et al., 2010). Such collective effort should be also implemented in the field of NFT to effectively evaluate its effects.

## Conclusions

The current meta-analysis showed significant effects of NFT for motor performance improvements after the last NFT session, but there were publication biases and substantial heterogeneity among the trials. Furthermore, there was a dose-response gradient between NFTs and motor performance improvements and reliable benefits when the NFT is performed longer than 1 week. The effectiveness of NFT for each motor performance measure remains unclear because of high heterogeneity or a small number of trials. The lasting effect after the final NFT session would also be a target for future research. Further accumulation of empirical NFT studies for motor performance improvement will be necessary to provide reliable evidence of the NFT effects on specific motor skills and safely incorporate NFT in the real world.

## Abbreviations

CI: confidence interval
EEG: electroencephalogram
fMRI: functional magnetic resonance imaging
fNIRS: functional near-infrared spectroscopy
MEG: magnetoencephalogram
NFT: neurofeedback training
NRS: non-randomized controlled study
RCT: randomized controlled trial
SMD: standardized mean difference

## Acknowledgements

This work was conducted as a part of Braintech Guidebook development in JST Moonshot R&D (Grant Number JPMJMS2012). We thank the members of the Evidence Evaluation Committee for Braintech Guidebook, a specially organized group for the Moonshot R&D project, for their comments on the manuscript. Special thanks also go to Prof. Takeo Nakayama and Dr. Yuko Masuzawa for their thoughtful advice on the process of systematic review and meta-analysis.

## References

Al-Wasity, S., Vogt, S., Vuckovic, A., Pollick, F.E., 2021. Upregulation of Supplementary Motor Area Activation with fMRI Neurofeedback during Motor Imagery. eNeuro 8, 1–14. https://doi.org/10.1523/ENEURO.0377-18.2020

Arns, M., de Ridder, S., Strehl, U., Breteler, M., Coenen, A., 2009. Efficacy of neurofeedback treatment in ADHD: the effects on inattention, impulsivity and hyperactivity: a meta-analysis. Clin. EEG Neurosci. 40, 180–189. https://doi.org/10.1177/155005940904000311

Arns, M., Kleinnijenhuis, M., Fallahpour, K., Breteler, R., 2008. Golf performance enhancement and real-life neurofeedback training using personalized event-locked EEG profiles. J. Neurother. 11, 11–18. https://doi.org/10.1080/10874200802149656

Bai, Z., Fong, K.N.K., Zhang, J.J., Chan, J., Ting, K.H., 2020. Immediate and long-term effects of BCI-based rehabilitation of the upper extremity after stroke: a systematic review and meta-analysis. J. Neuroeng. Rehabil. 17, 57. https://doi.org/10.1186/s12984-020-00686-2

Boe, S., Gionfriddo, A., Kraeutner, S., Tremblay, A., Little, G., Bardouille, T., 2014. Laterality of brain activity during motor imagery is modulated by the provision of source level neurofeedback. Neuroimage 101, 159–167. https://doi.org/10.1016/j.neuroimage.2014.06.066

Cheng, M.-Y., Huang, C.-J., Chang, Y.-K., Koester, D., Schack, T., Hung, T.-M., 2015. Sensorimotor Rhythm Neurofeedback Enhances Golf Putting Performance. J. Sport Exerc. Psychol. 37, 626–636. https://doi.org/10.1123/jsep.2015-0166

Cheng, M.-Y., Wang, K.-P., Hung, C.-L., Tu, Y.-L., Huang, C.-J., Koester, D., Schack, T., Hung, T.-M., 2017. Higher power of sensorimotor rhythm is associated with better performance in skilled air-pistol shooters. Psychol. Sport Exerc. 32, 47–53. https://doi.org/10.1016/j.psychsport.2017.05.007

Chiba, T., Kanazawa, T., Koizumi, A., Ide, K., Taschereau-Dumouchel, V., Boku, S., Hishimoto, A., Shirakawa, M., Sora, I., Lau, H., Yoneda, H., Kawato, M., 2019. Current Status of Neurofeedback for Post-traumatic Stress Disorder: A Systematic Review and the Possibility of Decoded Neurofeedback. Front. Hum. Neurosci. 13, 233. https://doi.org/10.3389/fnhum.2019.00233

Chiew, M., LaConte, S.M., Graham, S.J., 2012. Investigation of fMRI neurofeedback of differential primary motor cortex activity using kinesthetic motor imagery. Neuroimage 61, 21–31. https://doi.org/10.1016/j.neuroimage.2012.02.053

Christie, S., Bertollo, M., Werthner, P., 2020. The Effect of an Integrated Neurofeedback and Biofeedback Training Intervention on Ice Hockey Shooting Performance. J. Sport Exerc. Psychol. 42, 1–14. https://doi.org/10.1123/jsep.2018-0278

Cortese, S., Ferrin, M., Brandeis, D., Holtmann, M., Aggensteiner, P., Daley, D., Santosh, P., Simonoff, E., Stevenson, J., Stringaris, A., Sonuga-Barke, E.J.S., European ADHD Guidelines Group (EAGG), 2016. Neurofeedback for Attention-Deficit/Hyperactivity Disorder: Meta-Analysis of Clinical and Neuropsychological Outcomes From Randomized Controlled Trials. J. Am. Acad. Child Adolesc. Psychiatry 55, 444–455. https://doi.org/10.1016/j.jaac.2016.03.007

Domingos, C., Peralta, M., Prazeres, P., Nan, W., Rosa, A., Pereira, J.G., 2021. Session Frequency Matters in Neurofeedback Training of Athletes. Appl. Psychophysiol. Biofeedback 46, 195–204. https://doi.org/10.1007/s10484-021-09505-3

Doppelmayr, M., Weber, E., 2011. Effects of SMR and theta/beta neurofeedback on reaction times, spatial abilities, and creativity. J. Neurother. 15, 115–129. https://doi.org/10.1080/10874208.2011.570689

Egner, T., Gruzelier, J.H., 2004. EEG biofeedback of low beta band components: frequency-specific effects on variables of attention and event-related brain potentials. Clin. Neurophysiol. 115, 131–139. https://doi.org/10.1016/s1388-2457(03)00353-5

Enriquez-Geppert, S., Smit, D., Pimenta, M.G., Arns, M., 2019. Neurofeedback as a Treatment Intervention in ADHD: Current Evidence and Practice. Curr. Psychiatry Rep. 21, 46. https://doi.org/10.1007/s11920-019-1021-4

Fernández-Alvarez, J., Grassi, M., Colombo, D., Botella, C., Cipresso, P., Perna, G., Riva, G., 2022. Efficacy of bio- and neurofeedback for depression: a meta-analysis. Psychol. Med. 52, 201–216. https://doi.org/10.1017/S0033291721004396

Fujimoto, H., Mihara, M., Hattori, N., Hatakenaka, M., Yagura, H., Kawano, T., Miyai, I., Mochizuki, H., 2017. Neurofeedback-induced facilitation of the supplementary motor area affects postural stability. Neurophotonics 4, 045003. https://doi.org/10.1117/1.NPh.4.4.045003

Gong, A., Nan, W., Yin, E., Jiang, C., Fu, Y., 2020. Efficacy, Trainability, and Neuroplasticity of SMR vs. Alpha Rhythm Shooting Performance Neurofeedback Training. Front. Hum. Neurosci. 14, 94. https://doi.org/10.3389/fnhum.2020.00094

Gruzelier, J.H., Thompson, T., Redding, E., Brandt, R., Steffert, T., 2014. Application of alpha/theta neurofeedback and heart rate variability training to young contemporary dancers: state anxiety and creativity. Int. J. Psychophysiol. 93, 105–111. https://doi.org/10.1016/j.ijpsycho.2013.05.004

Hammond, D.C., 2011. Placebos and Neurofeedback: A Case for Facilitating and Maximizing Placebo Response in Neurofeedback Treatments. J. Neurother. 15, 94–114. https://doi.org/10.1080/10874208.2011.570694

He, S., Everest-Phillips, C., Clouter, A., Brown, P., Tan, H., 2020. Neurofeedback-Linked Suppression of Cortical β Bursts Speeds Up Movement Initiation in Healthy Motor Control: A Double-Blind Sham-Controlled Study. J. Neurosci. 40, 4021–4032. https://doi.org/10.1523/JNEUROSCI.0208-20.2020

Higgins, J.P.T., Altman, D.G., Gøtzsche, P.C., Jüni, P., Moher, D., Oxman, A.D., Savovic, J., Schulz, K.F., Weeks, L., Sterne, J.A.C., Cochrane Bias Methods Group, Cochrane Statistical Methods Group, 2011. The Cochrane Collaboration’s tool for assessing risk of bias in randomised trials. BMJ 343, d5928. https://doi.org/10.1136/bmj.d5928

Kajal, D.S., Braun, C., Mellinger, J., Sacchet, M.D., Ruiz, S., Fetz, E., Birbaumer, N., Sitaram, R., 2017. Learned control of inter-hemispheric connectivity: Effects on bimanual motor performance. Hum. Brain Mapp. 38, 4353–4369. https://doi.org/10.1002/hbm.23663

Kilavik, B.E., Zaepffel, M., Brovelli, A., MacKay, W.A., Riehle, A., 2013. The ups and downs of β oscillations in sensorimotor cortex. Exp. Neurol. 245, 15–26. https://doi.org/10.1016/j.expneurol.2012.09.014

Koizumi, A., Amano, K., Cortese, A., Shibata, K., Yoshida, W., Seymour, B., Kawato, M., Lau, H., 2016. Fear reduction without fear through reinforcement of neural activity that bypasses conscious exposure. Nature Human Behaviour 1, 1–7. https://doi.org/10.1038/s41562-016-0006

Landers, D.M., Petruzzello, S.J., Salazar, W., Crews, D.J., Kubitz, K.A., Gannon, T.L., Han, M., 1991. The influence of electrocortical biofeedback on performance in pre-elite archers. Med. Sci. Sports Exerc. 23, 123–129. https://doi.org/10.1249/00005768-199101000-00018

Little, S., Bonaiuto, J., Barnes, G., Bestmann, S., 2019. Human motor cortical beta bursts relate to movement planning and response errors. PLoS Biol. 17, e3000479. https://doi.org/10.1371/journal.pbio.3000479

Lofthouse, N., Arnold, L.E., Hersch, S., Hurt, E., DeBeus, R., 2012. A review of neurofeedback treatment for pediatric ADHD. J. Atten. Disord. 16, 351–372. https://doi.org/10.1177/1087054711427530

Luctkar-Flude, M., Groll, D., 2015. A Systematic Review of the Safety and Effect of Neurofeedback on Fatigue and Cognition. Integr. Cancer Ther. 14, 318–340. https://doi.org/10.1177/1534735415572886

Marins, T., Rodrigues, E.C., Bortolini, T., Melo, B., Moll, J., Tovar-Moll, F., 2019. Structural and functional connectivity changes in response to short-term neurofeedback training with motor imagery. Neuroimage 194, 283–290. https://doi.org/10.1016/j.neuroimage.2019.03.027

Maszczyk, A., Gołaś, A., Pietraszewski, P., Kowalczyk, M., Cięszczyk, P., Kochanowicz, A., Smółka, W., Zając, A., 2018. Neurofeedback for the enhancement of dynamic balance of judokas. Biol. Sport 35, 99–102. https://doi.org/10.5114/biolsport.2018.71488

Micoulaud-Franchi, J.-A., Geoffroy, P.A., Fond, G., Lopez, R., Bioulac, S., Philip, P., 2014. EEG neurofeedback treatments in children with ADHD: an updated meta-analysis of randomized controlled trials. Front. Hum. Neurosci. 8, 906. https://doi.org/10.3389/fnhum.2014.00906

Mikicin, M., Orzechowski, G., Jurewicz, K., Paluch, K., Kowalczyk, M., Wróbel, A., 2015. Brain-training for physical performance: a study of EEG-neurofeedback and alpha relaxation training in athletes. Acta Neurobiol. Exp. 75, 434–445.

Minds Manual Developing Committee (Ed.), 2021. Minds Manual for Guideline Development 2020 ver. 3.0. Japan Council for Quality Health Care, Tokyo.

Mirifar, A., Beckmann, J., Ehrlenspiel, F., 2017. Neurofeedback as supplementary training for optimizing athletes’ performance: A systematic review with implications for future research. Neurosci. Biobehav. Rev. 75, 419–432. https://doi.org/10.1016/j.neubiorev.2017.02.005

Mottola, F., Blanchfield, A., Hardy, J., Cooke, A., 2021. EEG neurofeedback improves cycling time to exhaustion. Psychol. Sport Exerc. 55, 101944. https://doi.org/10.1016/j.psychsport.2021.101944

Nan, W., Yang, L., Wan, F., Zhu, F., Hu, Y., 2020. Alpha down-regulation neurofeedback training effects on implicit motor learning and consolidation. J. Neural Eng. 17, 026014. https://doi.org/10.1088/1741-2552/ab7c1b

Ota, Y., Takamoto, K., Urakawa, S., Nishimaru, H., Matsumoto, J., Takamura, Y., Mihara, M., Ono, T., Nishijo, H., 2020. Motor Imagery Training With Neurofeedback From the Frontal Pole Facilitated Sensorimotor Cortical Activity and Improved Hand Dexterity. Front. Neurosci. 14, 34. https://doi.org/10.3389/fnins.2020.00034

Paul, M., Ganesan, S., Sandhu, J.S., Simon, J.V., 2012. Effect of sensory motor rhythm neurofeedback on psycho-physiological, electro-encephalographic measures and performance of archery players. Ibnosina Journal of Medicine and Biomedical Sciences 4, 32. https://doi.org/10.4103/1947-489X.210753

Raymond, J., Sajid, I., Parkinson, L.A., Gruzelier, J.H., 2005. Biofeedback and dance performance: a preliminary investigation. Appl. Psychophysiol. Biofeedback 30, 64–73. https://doi.org/10.1007/s10484-005-2175-x

Reiner, M., Lev, D.D., Rosen, A., 2018. Theta Neurofeedback Effects on Motor Memory Consolidation and Performance Accuracy: An Apparent Paradox? Neuroscience 378, 198–210. https://doi.org/10.1016/j.neuroscience.2017.07.022

Ring, C., Cooke, A., Kavussanu, M., McIntyre, D., Masters, R., 2015. Investigating the efficacy of neurofeedback training for expediting expertise and excellence in sport. Psychol. Sport Exerc. 16, 118–127. https://doi.org/10.1016/j.psychsport.2014.08.005

Rogel, A., Guez, J., Getter, N., Keha, E., Cohen, T., Amor, T., Todder, D., 2015. Transient Adverse Side Effects During Neurofeedback Training: A Randomized, Sham-Controlled, Double Blind Study. Appl. Psychophysiol. Biofeedback 40, 209–218. https://doi.org/10.1007/s10484-015-9289-6

Rostami, R., Sadeghi, H., Karami, K.A., Abadi, M.N., Salamati, P., 2012. The effects of neurofeedback on the improvement of rifle shooters’ performance. J. Neurother. 16, 264–269. https://doi.org/10.1080/10874208.2012.730388

Ros, T., Munneke, M.A.M., Parkinson, L.A., Gruzelier, J.H., 2014. Neurofeedback facilitation of implicit motor learning. Biol. Psychol. 95, 54–58. https://doi.org/10.1016/j.biopsycho.2013.04.013

Rozengurt, R., Barnea, A., Uchida, S., Levy, D.A., 2016. Theta EEG neurofeedback benefits early consolidation of motor sequence learning. Psychophysiology 53, 965–973. https://doi.org/10.1111/psyp.12656

Schulz, K.F., Altman, D.G., Moher, D., CONSORT Group, 2010. CONSORT 2010 statement: updated guidelines for reporting parallel group randomized trials. Obstet. Gynecol. 115, 1063–1070. https://doi.org/10.1097/AOG.0b013e3181d9d421

Senkowski, D., Molholm, S., Gomez-Ramirez, M., Foxe, J.J., 2006. Oscillatory beta activity predicts response speed during a multisensory audiovisual reaction time task: a high-density electrical mapping study. Cereb. Cortex 16, 1556–1565. https://doi.org/10.1093/cercor/bhj091

Sherlin, L.H., Ford, N.C.L., Baker, A.R., Troesch, J., 2015. Observational Report of the Effects of Performance Brain Training in Collegiate Golfers. Biofeedback Self. Regul. 43, 64–72. https://doi.org/10.5298/1081-5937-43.2.06

Shibata, K., Watanabe, T., Sasaki, Y., Kawato, M., 2011. Perceptual learning incepted by decoded fMRI neurofeedback without stimulus presentation. Science 334, 1413–1415. https://doi.org/10.1126/science.1212003

Sidhu, A., Cooke, A., 2021. Electroencephalographic neurofeedback training can decrease conscious motor control and increase single and dual-task psychomotor performance. Exp. Brain Res. 239, 301–313. https://doi.org/10.1007/s00221-020-05935-3

Strizhkova, O., Cherapkina, L., Strizhkova, T., 2012. Neurofeedback course applying of high skilled gymnasts in competitive period. J. Hum. Sport Exerc. 7, S185–S193. https://doi.org/10.4100/jhse.2012.7.proc1.21

Trambaiolli, L.R., Kohl, S.H., Linden, D.E.J., Mehler, D.M.A., 2021. Neurofeedback training in major depressive disorder: A systematic review of clinical efficacy, study quality and reporting practices. Neurosci. Biobehav. Rev. 125, 33–56. https://doi.org/10.1016/j.neubiorev.2021.02.015

Tzagarakis, C., Ince, N.F., Leuthold, A.C., Pellizzer, G., 2010. Beta-band activity during motor planning reflects response uncertainty. J. Neurosci. 30, 11270–11277. https://doi.org/10.1523/JNEUROSCI.6026-09.2010

Umeno, K., Nakamura, K., Inomoto, A., Miyata, H., 2018. Examining the Effect of Simple Electroencephalograph Neurofeedback on Mental Practice. Rigakuryoho Kagaku. https://doi.org/10.1589/rika.33.901

van der Kolk, B.A., Hodgdon, H., Gapen, M., Musicaro, R., Suvak, M.K., Hamlin, E., Spinazzola, J., 2016. A Randomized Controlled Study of Neurofeedback for Chronic PTSD. PLoS One 11, e0166752. https://doi.org/10.1371/journal.pone.0166752

Viviani, G., Vallesi, A., 2021. EEG-neurofeedback and executive function enhancement in healthy adults: A systematic review. Psychophysiology 58, e13874. https://doi.org/10.1111/psyp.13874

Wang, T., Mantini, D., Gillebert, C.R., 2018. The potential of real-time fMRI neurofeedback for stroke rehabilitation: A systematic review. Cortex 107, 148–165. https://doi.org/10.1016/j.cortex.2017.09.006

Xiang, M.-Q., Hou, X.-H., Liao, B.-G., Liao, J.-W., Hu, M., 2018. The effect of neurofeedback training for sport performance in athletes: A meta-analysis. Psychol. Sport Exerc. 36, 114–122. https://doi.org/10.1016/j.psychsport.2018.02.004

Yakovlev, L., Syrov, N., Görtz, N., Kaplan, A., 2020. BCI-controlled motor imagery training can improve performance in e-sports, in: Communications in Computer and Information Science, Communications in Computer and Information Science. Springer International Publishing, Cham, pp. 581–586. https://doi.org/10.1007/978-3-030-50726-8_76

Yeh, W.-H., Hsueh, J.-J., Shaw, F.-Z., 2020. Neurofeedback of Alpha Activity on Memory in Healthy Participants: A Systematic Review and Meta-Analysis. Front. Hum. Neurosci. 14, 562360. https://doi.org/10.3389/fnhum.2020.562360

